# Defining a critical role of an essential membrane protein in mycolic acid transport in mycobacteria

**DOI:** 10.1101/2023.05.02.539095

**Authors:** Jeremy Liang, Yushu Chen, Shu-Sin Chng

**Author notes:** To whom correspondences should be addressed: Yushu Chen,; Shu-Sin Chng,. These authors contributed equally to this work (Jeremy Liang, Yushu Chen).

## Abstract

A major feature of the mycobacterial outer membrane (OM) is the presence of long, branched chain mycolic acids (MAs), which render the OM hydrophobic and impervious against various noxious substances, including antibiotics. While the biosynthesis of MA is well studied, the mechanisms governing its transport from the inner membrane to the OM remain largely elusive. In this study, we characterized the function of MSMEG_0317 in *Mycobacterium smegmatis*, a membrane protein encoded within a conserved genetic locus that has been implicated in MA metabolism and/or transport. Using a conditional knockout mutant, we demonstrate that *msmeg_0317* is essential for mycobacterial growth. Depleting *msmeg_0317* from cells blocks the formation of MA species found at the OM, establishing a critical function in MA transport across the cell envelope. We further reveal that MSMEG_0317 exists as stable dimers *in vitro* that require the presence of its N- and C-terminal transmembrane helices, both of which are important for functionality in cells. Our work defines the essential role of MSMEG_0317 in MA metabolism and/or transport, and offers new insights into cell envelope biogenesis in mycobacteria.

## Introduction

The outer membrane (OM) of *Mycobacterium tuberculosis*, the causative agent of tuberculosis (TB), is distinctively enriched in C_60_–C_90_ long-chain branched mycolic acids (MAs) that are packed together with other lipids to produce a highly hydrophobic and impervious bilayer (1). This impermeability in part confers mycobacterial cells with intrinsic resistance against many antibiotics. In addition, MAs are essential for the growth and survival of all mycobacterial species. In the OM, these MAs comprise extractable mycolates, such as trehalose monomycolate (TMM) and dimycolate (TDM), and non-extractable mycolates that are covalently bound to arabinogalactan polysaccharides, in turn linked to the peptidoglycan, and collectively known as the mAGP complex (2,3). MAs are first biosynthesized as TMMs at the inner membrane (IM) via a highly conserved and well-characterized pathway, which is the target for first-line anti-TB drug isoniazid (4). Newly synthesized TMMs are flipped across the IM by Mycobacterial membrane protein Large 3 (MmpL3), and subsequently transported to the OM via an unknown mechanism (5). There, the single mycolate chain on TMM is transferred to another TMM molecule to form TDM, or to the arabinogalactan polysaccharide to form the mAGP complex, processes mediated by the antigen-85 (Ag85) enzymes/complex (6).

Significant gaps remain in our detailed understanding of how TMMs are flipped across the IM, and/or are delivered across the periplasmic space. Recent studies in *Corynebacterium glutamicum* suggest that several proteins with yet to be characterized function(s) may modulate these TMM transport processes, either directly or indirectly (7-9). *C. glutamicum* is a related species in the *Mycobacteriales* order that produces shorter C_22_–C_36_ versions of MAs known as corynomycolic acids (CMAs), which are non-essential (10). Notably, it has been shown that trehalose monocorynomycolate (TMCM) may be modified before/during transport by a putative acetyltransferase TmaT (9,11). Deletion of *tmaT* resulted in an accumulation of TMCM, and the loss of both trehalose dicorynomycolate (TDCM) (9), a phenomenon also observed with TMM/TDM in an *M. smegmatis* conditional *tmaT* knockout mutant (12). Furthermore, *C. glutamicum* cells lacking TmaT no longer produce a distinct acetylated-TMCM species, thus proposed to be an intermediate in TMCM transport (9). Interestingly, *tmaT* is co-located at a conserved locus with other genes (*mtrP* and *mmpA*) now shown to be implicated in TMCM acetylation and/or transport, even though similar phenotypes are not yet demonstrated in mycobacteria (Fig. 1A) (7,8). How these genes/proteins, and potentially others found/encoded in the same locus, influence TM(C)M metabolism and transport are still unclear.

**Figure 1:**
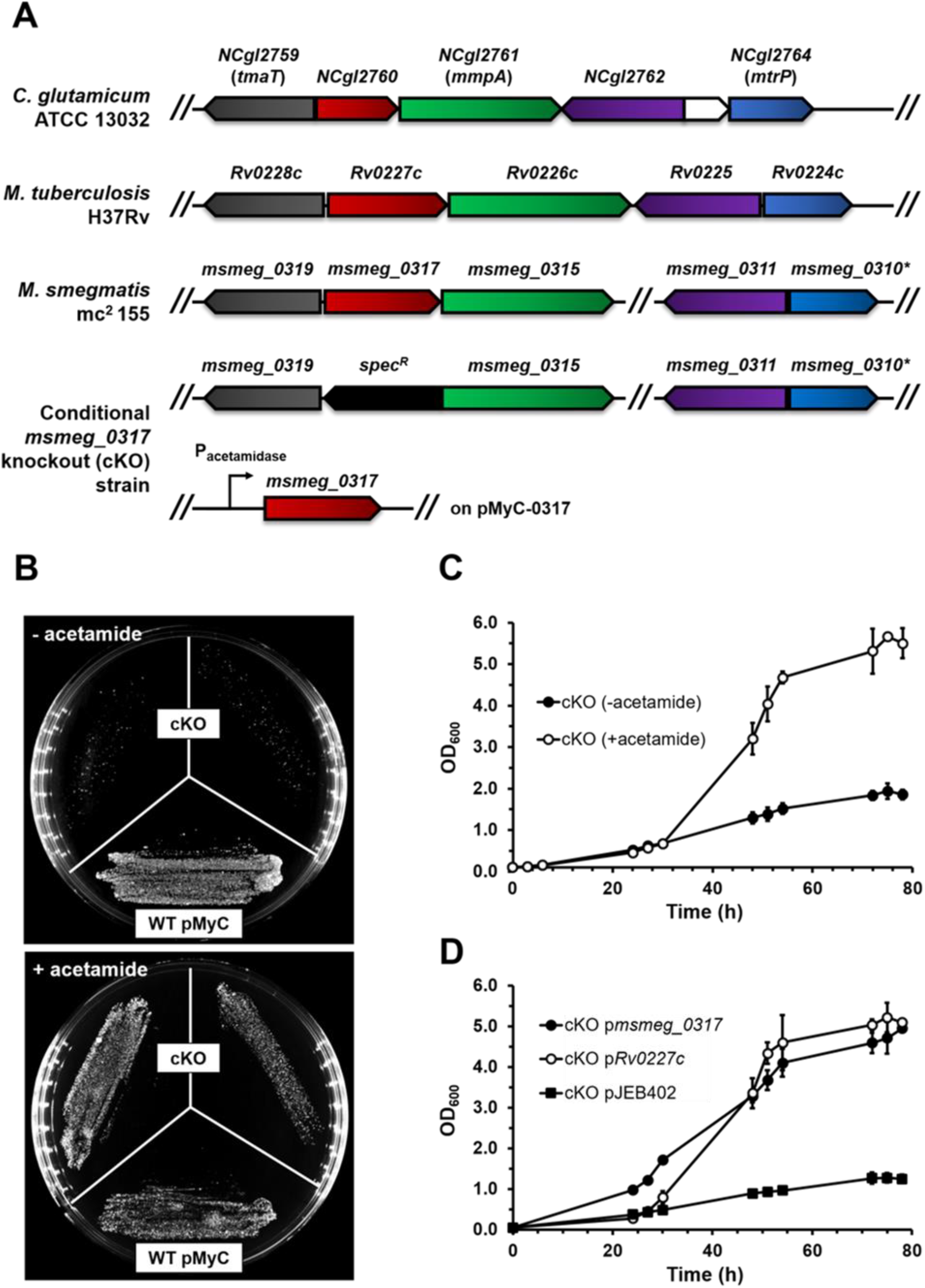
*msmeg_0317* is a conserved essential gene. **(A)** (*top*) Organization of the genetic locus of *msmeg_0317* and its orthologs in *C. glutamicum* and *M. tuberculosis*. Orthologous genes are represented by the same colors; asterisks denote an annotated pseudogene. (*bottom*) Genotype of the conditional *msmeg_0317* knockout (cKO) strain. **(B)** Streaked cultures of WT harboring empty vector (pMyC) and the cKO strain on LB solid medium in the presence or absence of acetamide. **(C)** Growth curves of the cKO strain in liquid 7H9 medium in the presence or absence of acetamide. **(D)** Growth curves of cKO cells constitutively expressing *msmeg_0317* or *rv0227c* from the integrative pJEB402 plasmid in liquid 7H9 medium in the absence of acetamide. Error bars represent standard deviation from biological triplicates.

Another gene of particular interest in the *tmaT* locus is *NCgl2760* or *Rv0227c* in *C. glutamicum* or *M. tuberculosis*, respectively. NCgl2760 has been suggested to be involved in the elongation of lipoglycans, specifically lipomannans (LM) and lipoarabinomannans (LAM) (13). However, Rv0227c appears to be essential for mycobacterial growth even though LM/LAM are not (14-16), indicating that biosynthesis of these lipoglycans might not be its key physiological function in mycobacteria. Curiously, Rv0227c has been predicted by phylogenetic profiling to be functionally linked to MA biosynthesis (17). Therefore, whether and/or how Rv0227c and its homologs are involved in TMM metabolism and transport in mycobacteria warrant further investigation. Here, we establish that MSMEG_0317 is indeed required for TMM transport in *M. smegmatis*. We verify that MSMEG_0317 is essential for growth. In addition, we show that MSMEG_0317 exists as stable dimers, and dimerization may be required for its function in cells. Importantly, we reveal that depletion of MSMEG_0317 causes significant reductions in TDM and mAGP formation, implicating this protein in TMM transport across the cell envelope. Our work adds to and highlights the strong connection between the *tmaT*-*Rv0227c-mmpA-Rv0225*-*mtrP* genetic locus and MA metabolism and transport.

## Results

### *msmeg_0317* is a conserved gene essential for mycobacterial growth

Multiple TRADIS experiments have indicated that *Rv0227c* is essential in *M. tuberculosis* (14,15). It has also been suggested that *msmeg_0317* is required for growth in *M. smegmatis*, owing to the inability to generate a clean deletion (13). To verify its essentiality and study its function, we generated a conditional *msmeg_0317* knockout (cKO) mutant in *M. smegmatis*. In this strain, chromosomal *msmeg_0317* was deleted in the presence of a second copy on an episomal plasmid under the control of an acetamide-inducible promoter (pMyC-0317). When supplemented with acetamide, the cKO mutant grew like WT cells, albeit with a longer lag phase. Consistent with the essentiality of *msmeg_0317*, the cKO mutant was unable to grow in the absence of acetamide (i.e., *msmeg_0317* depleted), either on solid or in liquid media (Fig. 1B & C). We further demonstrated that constitutive expression of *msmeg_0317* or *Rv0227c* can rescue growth of the cKO strain without acetamide (Fig. 1D). We conclude that *msmeg_0317* homologs are essential in mycobacteria with conserved function(s).

### MSMEG_0317 likely functions as a dimeric protein

To gain insights into the function(s) of MSMEG_0317, we attempted to identify potential interacting partners via native affinity purification (AP). MmpL3 and TmaT have been suggested as interacting partners based on bacterial two-hybrid screens (18); however, we did not identify significant protein hits enriched by C-terminally His-tagged MSMEG_0317 expressed from an integrative plasmid and purified from *M. smegmatis* cells. To further characterize the protein, we overexpressed the same His-tagged construct in *Escherichia coli* and purified it to homogeneity under non-denaturing conditions. We showed that MSMEG_0317 eluted as a single peak on size exclusion chromatography (SEC), much earlier than expected for a 44-kDa protein (Fig. 2A). Multi-angle light scattering (MALS) analysis revealed the estimated molecular weight of this peak to be ∼78.7 (± 1.6%) kDa (Fig. 2B), indicating the presence of a stable dimeric species (expected size 88.2 kDa). We further observed that purified MSMEG_0317 exhibited anomalous migration (smear ∼35-90 kDa) on SDS-PAGE when samples were not heated, a phenomenon that may be related to its oligomeric state, particularly as a membrane protein (Fig. 2A). Our results indicate that MSMEG_0317 may function as dimers.

**Figure 2:**
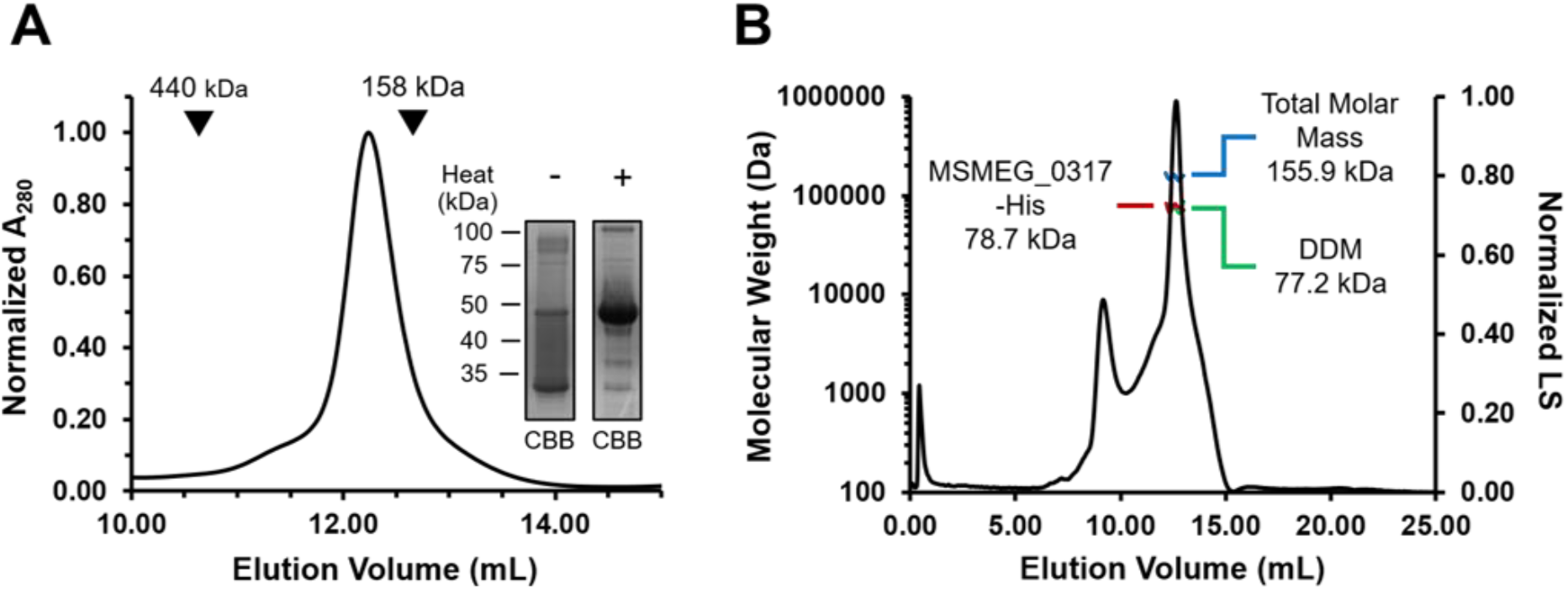
MSMEG_0317 forms stable dimers *in vitro*. **(A)** SEC profile of MSMEG_0317-His. Elution volume of molecular weight markers (440 kDa & 158 kDa) are annotated with arrow heads. Inset shows the heat-modifiable gel shift exhibited by purified MSMEG_0317-His on SDS-PAGE. **(B)** SEC-MALS analysis of MSMEG_0317-His. Total molar mass: 155.9 (±1.6%) kDa; modifier (DDM) molar mass: 77.2 (±3.6%); protein molar mass: 78.7 (±1.6%) kDa (measured), 88.2 kDa (predicted for MSMEG_0317-His dimer). LS, light scattering.

MSMEG_0317 is a membrane protein predicted to contain two transmembrane helices flanking a PorA periplasmic domain (296 aa), accompanied by a C-terminal cytoplasmic domain (50 aa) (Fig. 3A). To identify structural elements required for MSMEG_0317 dimerization and/or function, we constructed, purified, and characterized several truncated variants that lack either the N-terminal TM helix (0317ΔNH), C-terminal TM helix (0317ΔCHD), the cytoplasmic domain (0317ΔCD), or all of these elements (S-0317) (Fig. 3B). We also tested if these truncated variants could support the growth of the *msmeg_0317* cKO mutant. The only variant that exhibits stable dimerization is 0317ΔCD, as judged by SEC-MALS analysis (83.2 (± 4.4%) kDa; expected size 72.5 kDa) (Fig. 3C & S1A). In addition, introducing *0317ΔCD* into the cKO mutant can fully complement growth in the absence of acetamide (Fig. 3D). Therefore, the cytoplasmic domain is not required for protein dimerization and function. 0317ΔCHD underwent degradation *in vitro* and was excluded from further analysis. We noted that 0317ΔNH can still form dimers (MALS: 78.3 (± 4.4 %) kDa; expected size 82.6 kDa) but tends to aggregate on SEC (Fig. S1B), suggesting instability. Both 0317ΔCHD and 0317ΔNH were unable to rescue the growth of the cKO mutant with no acetamide (Fig. 3C), despite being expressed at levels comparable to 0317ΔCD (Fig. S2). Interestingly, we found that S-0317 forms only monomers on SEC (MALS: 37.7 (± 8.6%) kDa; expected size 33.5 kDa) (Fig. S1C); we attempted to target S-0317 to the periplasm using appropriate signal sequences, but these were somehow not processed in cells. Taken together, our data reveal that both transmembrane helices appear to be important for MSMEG_0317 stability and/or dimerization, and are required for physiological function.

**Figure 3:**
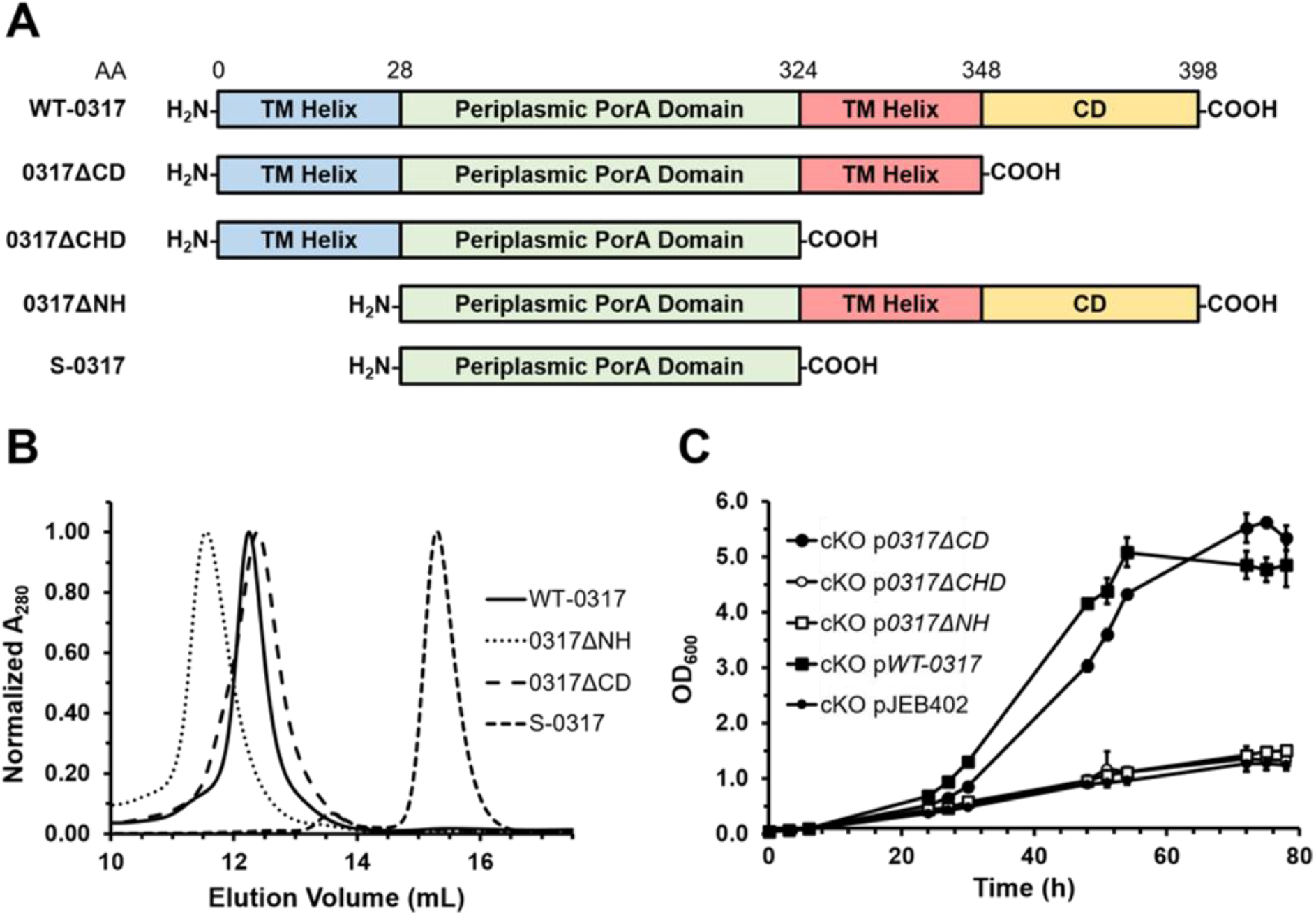
Both TM helices of MSMEG_0317 are important for dimerization and physiological function. **(A)** Illustration of the predicted structural elements of WT MSMEG_0317 (WT-0317) and the truncated MSMEG_0317 mutants generated. **(B)** SEC profile of the WT and truncated MSMEG_0317 proteins. 0317ΔCHD is not shown since it degraded during SEC purification. **(C)** Growth curves of cKO cells constitutively expressing WT or the truncated *msmeg_0317* variants from the integrative pJEB402 plasmid in liquid 7H9 medium in the absence of acetamide. Error bars represent standard deviation from biological triplicates.

### MSMEG_0317 is required for TMM transport to the OM

In mycobacteria, *msmeg_0317* and its homologs are found in a genetic locus where many genes have been implicated in MA metabolism (7-9). Given its essentiality, we sought to elucidate whether *msmeg_0317* also has functional roles in related processes. We profiled extractable and non-extractable lipids newly synthesized in the *msmeg_0317* cKO strain via [^14^C]-acetate labelling of cells grown with or without acetamide. Even though the levels of TMM remained unchanged, thin-layer chromatography (TLC) analyses revealed that there was essentially no or little newly-synthesized TDM and mAGP in the *msmeg_0317*-depleted cKO cells, compared to the non-depleted cells (Fig. 4A & B). In addition, we observed the appearance of two unknown lipid spots on TLC, one of which migrated right above TMM, perhaps implying structural similarity to the latter (Fig. 4A). We conclude that MSMEG_0317 is required for TMM transport across the mycobacterial cell envelope.

**Figure 4:**
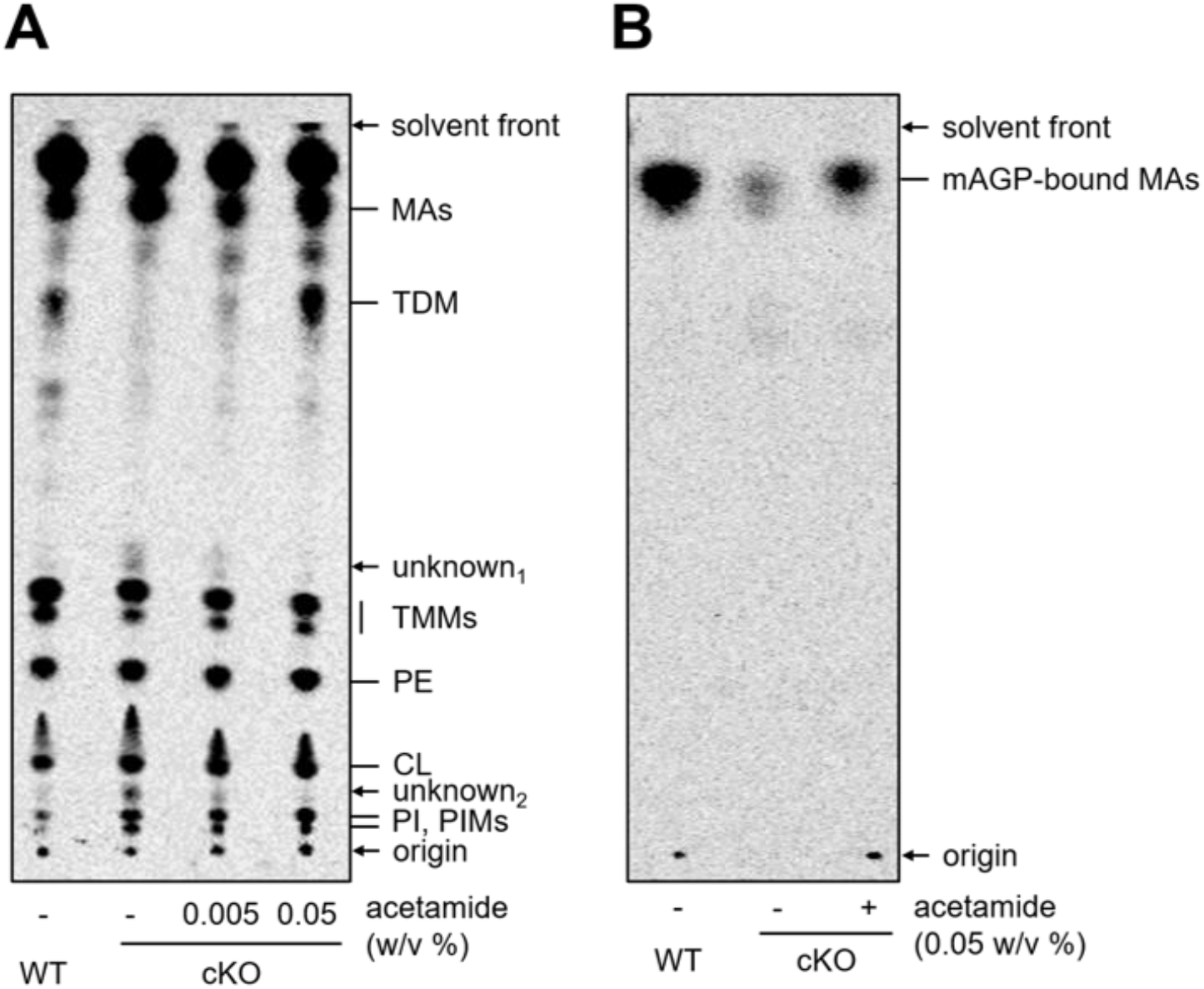
MSMEG_0317 is involved in TMM transport in mycobacteria. TLC analysis of [^14^C]-labelled **(A)** total extractable lipids and **(B)** arabinogalactan-bound MAs extracted from WT cells and cKO strains, grown in the presence of different concentrations of acetamide, visualized by phosphor imaging. TLCs were developed with a solvent system comprising chloroform-methanol-water (30:8:1). Lipids were assigned based on previously reported retention factors obtained in the same solvent system via TLC (3). CL, cardiolipin; PE, phosphatidylethanolamine; PI, phosphatidylinositol; PIM, phosphatidylinositol mannoside.

## Discussion

In this work, we probed the function of MSMEG_0317 and the biochemical basis thereof. We have shown that *msmeg_0317* is an essential gene in mycobacteria required for TMM transport to the OM. We have also demonstrated that MSMEG_0317 forms stable dimers *in vitro*; both predicted TM helices of the protein appear to be important for its stability and function, while the C-terminal cytoplasmic domain is dispensable. Our study highlights a novel player in mycobacterial OM biogenesis.

How MSMEG_0317 may be involved in TMM metabolism and/or transport is not known. MSMEG_0317 is predicted to contain a PorA domain in the periplasm. Interestingly, the crystal structure of this domain reveals the presence of hydrophobic cavities that may have roles in lipid binding (19). Consistent with this idea, MSMEG_0317 has been preliminarily identified as a possible TMM-binding protein via an affinity-based protein profiling approach that utilized a photoactivatable C15-TMM analog (20). While the validity and/or specificity of proposed MSMEG_0317-TMM interactions remain to be ascertained, these studies suggest that MSMEG_0317 plays a direct role in TMM metabolism and/or transport. One possibility is that MSMEG_0317 may work with the TMM flippase MmpL3 to facilitate transport to the OM. MmpL3 contains a large periplasmic domain that can bind TMM, and may additionally be involved in TMM release from the IM (21,22); MSMEG_0317 could therefore function to usher TMM from MmpL3 to subsequent factors that shuttle TMM to the OM. Another possibility is that MSMEG_0317 could be involved in TMM modification steps thought to be important during TMM transport. If TmaT acetylation of TMM indeed occurs in mycobacteria, we speculate MSMEG_0317 may work with other protein(s) to deacetylate these modified TMMs at the periplasmic leaflet of the IM. In both of these scenarios, one might expect MSMEG_0317 to interact with either MmpL3 or TmaT, which has been previously suggested (18). However, we have not been able to detect any stable/specific interactions between MSMEG_0317 and either protein *in vitro*. Conversely, MSMEG_0317 may indirectly affect TMM metabolism and/or transport, where the protein somehow regulates this (and other) envelope biogenesis processes, possibly by binding TMMs at the IM. Such function(s) may explain why removal of MSMEG_0317 also apparently affects LM/LAM biosynthesis (13). We note that a very recent report proposed a possible role for MSMEG_0317 in coordinating processes important for polar growth in mycobacteria (23). The role of MSMEG_0317 in TMM transport and/or envelope biogenesis requires further investigations.

As a leading cause of death by infectious disease, TB remains a major global health threat to be tackled (24). Moreover, the emergence of multi-drug resistant TB infections presses the need for the discovery of new anti-mycobacterial drugs. In this regard, the essential MSMEG_0317/Rv0227c proteins represent ideal targets for drug inhibition. Furthermore, the genetic locus of *msmeg_0317*/*Rv0227c*, is home to several essential genes that have been implicated in TMM metabolism and/or transport. Elucidating the functions of the genes in this cluster would additionally yield potential protein targets for drug development.

## Experimental procedures

### Bacterial strains and growth conditions

*M. smegmatis* mc^2^155 was used for all *M. smegmatis* experiments. The *msmeg_0317* cKO strain was generated from wild-type *M. smegmatis* mc^2^155 and used in this study. *M. smegmatis* mc^2^155 was grown in 7H9 broth (BD Falcon) supplemented with 0.05% tyloxapol and 0.5% glycerol, or tryptic soy broth (TSB; BD Falcon) supplemented with 0.05% Tween 80, where specified, or on LB solid agar. The concentrations of antibiotics used for *M. smegmatis* are as follows: 20 μg/ml kanamycin (Kan; Sigma- Aldrich), 20 μg/ml spectinomycin (Spec; Sigma-Aldrich), 50 μg/ml hygromycin (Hyg; Merck Millipore). *E. coli* NovaBlue and BL21(λDE3) cells were used for cloning purposes and protein expression respectively, and were grown in LB broth or on LB solid agar at 37 °C. The concentrations of antibiotics used for *E. coli* are as follows: 50 μg/ml Kan, 50 μg/ml Spec, 150 μg/ml Hyg, 100 μg/ml ampicillin (Amp; Sigma-Aldrich).

### Plasmid construction

To construct most plasmids, the genes-of-interest or DNA fragments were amplified by PCR from their DNA templates with the appropriate primers. The amplified fragments were either digested with relevant restriction enzymes (New England Biolabs) and ligated into the same sites on their stipulated plasmids using T4 DNA ligase (New England Biolabs), or assembled with other fragments via Gibson assembly with the ClonExpress Ultra One Step Cloning Kit (Vazyme Biotech). *E. coli* NovaBlue competent cells were then transformed with the ligation/assembled products and selected on LB plates containing appropriate antibiotics. All constructs were verified by DNA sequencing (Bio Basic Asia Pacific, Singapore). All plasmids and primers used can be found in Table S1 and S2 respectively.

### Construction of *msmeg*_*0317* cKO strain and derivative strains

The *msmeg_0317* cKO strain was constructed *via* two-step homologous recombination as previously described (25). Briefly, a pYUB854 vector (Hyg^R^) was constructed bearing a *spec*^*R*^ cassette flanked by the 5’- and 3’-untranslated regions of the *msmeg_0317* gene, a counter-selectable marker *sacB* and a reporter gene *lacZ* and. The suicide plasmid was electroporated into WT *M. smegmatis* cells and was integrated into the genome at the native locus of *msmeg_0317* via homologous recombination. Successful first cross-over transformants (blue colonies) were selected and screened on Hyg/Spec agar plates supplemented with 20 μg/ml X-gal (Bio Basic Asia Pacific, Singapore). A Kan^R^ episomal plasmid containing *msmeg_0317* under the control of an acetamide-inducible promoter (pMyC-0317) was transformed into the first cross-over cells via electroporation. The resulting cells were then plated on Spec/Kan agar plates supplemented with 5% sucrose and 0.05% acetamide (Sigma-Aldrich), to induce *msmeg_0317* deletion via the second crossover. The cKO strain generated was verified via colony PCR and DNA sequencing. For complementation studies, *Rv0227c* and the truncated *msmeg_0317* variant genes were cloned into a Hyg^R^ pJEB402 integrative plasmid, under the constitutive MOP promoter. For the native affinity purification of MSMEG_0317, a copy of *msmeg_0317* fused with either N- or C-terminal His-tag coding sequence was cloned into a Kan^R^ pJEB402 plasmid under the MOP constitutive promoter.

### Growth measurement

The cKO strain and its derivatives were inoculated and grown till the log phase in 7H9 media supplemented with 0.05% acetamide. To deplete *msmeg_0317*, cells were washed twice with 7H9 media and sub-cultured into fresh media supplemented with or without acetamide. Growth curves were generated across 78 hours of growth, with triplicate samples taken at stipulated time intervals. For plate streaking experiments, washed cells were spotted onto LB agar and streaked with an inoculation loop.

### Native affinity purification of MSMEG_0317

*M. smegmatis* cells expressing His-tagged MSMEG_0317 from the appropriate plasmids were grown in 7.5 mL of TSB to stationary phase (OD_600_ ∼5) and subcultured into 750 mL of TSB. This culture was grown to stationary phase (OD_600_ ∼5) and harvested by centrifugation at 4,700 x *g* for 15 minutes. Harvested cells were resuspended in 12.5 mL of cold Tris buffered saline (TBS; 20 mM Tris-HCl pH 8.0, 150 mM NaCl) containing 1 mM phenylmethylsulfonyl fluoride (PMSF), 100 μg/mL lysozyme, and 50 μg/mL DNase I. The resuspended cells were lysed by two passages through a high-pressure French Press (GlenMills) homogenizer at 20,000 psi, after which unbroken cells were removed by centrifugation at 4,800 x *g* for 5 minutes. Membrane fractions were isolated from the cell lysate by ultracentrifugation (Model XL-90, Beckman Coulter) at 95,000 x *g* for 1 hour. Membrane proteins were extracted from the pellet with 7.5 mL of TBS-A (20 mM Tris-HCl pH 8.0, 150 mM NaCl, 5 mM MgCl_2_, 10% glycerol, 5 mM imidazole, and 1% n-dodecyl β-d-maltoside (DDM; Calbiochem) for 2 hours. This suspension was further subjected to ultracentrifugation at 89,000 x g for 1 hour. The supernatant was incubated with 125 μL of TALON cobalt resin (Takara Bio; pre-incubated with 5 mL of TBS supplemented with 5 mM of imidazole) for 2 hours at 4 °C. Resin suspension was loaded onto a column and the first flowthrough was recycled into the packed resin once more. Resins were washed 10 times with 500 μL of TBS-B (20 mM Tris-HCl pH 8.0, 300 mM NaCl, 10 mM imidazole, 5 mM MgCl_2_, 10% glycerol, 0.05% DDM), and eluted with 100 μL of TBS- C (20 mM Tris-HCl pH 8.0, 150 mM NaCl, 200 mM imidazole, 5 mM MgCl_2_, 10% glycerol, and 0.05% DDM). The eluate was concentrated with an Amicon Ultra 10-kDa cut-off ultrafiltration device (Merck Millipore) and subjected to SDS-PAGE analysis. Protein gel sections and bands to be identified were sent to Taplin Biological Mass Spectrometry Facility, Harvard Medical School (Boston, MA) for capillary LC-MS/MS analysis.

### Purification of MSMEG_0317 and truncated variants from *E. coli*

All proteins were overexpressed in and purified from respective transformed BL21(λDE3) strains. An overnight 7.5 mL culture was grown from a single colony in LB broth supplemented with appropriate antibiotics at 37 °C. This culture was then used to inoculate a 750 mL culture and grown at the same temperature until an OD_600_ of ∼0.5. For induction, 1 mM isopropyl β-d-thiogalactopyranoside (IPTG; Axil Scientific) was added, and the culture was grown for another 3 hours at the same temperature. Cells were harvested by centrifugation at 4,800 x *g* for 10 minutes, and resuspended in cold TBS. The same procedures from the native affinity purification of MSMEG_0317 were carried out to obtain affinity-purified recombinant His-tagged proteins. The eluate was further purified by SEC system (AKTA Pure; GE Healthcare) at 4 °C on a prepacked Superdex 200 increase 10/300 GL column, with TBS-D (20 mM Tris-HCl (pH 8.0) 150 mM NaCl, 10% glycerol, 5 mM MgCl_2_, and 0.05% DDM) as the eluent. Fractions corresponding to the peaks of interest were pooled and concentrated, before being subjected to SDS-PAGE and SEC-MALS analysis.

### Protein expression level testing in the cKO strain

cKO cells expressing His-tagged MSMEG_0317 or its truncated variants from appropriate plasmids were grown in 7H9 media supplemented with 0.05% acetamide until the stationary phase (OD_600_ ∼5). Cells were normalized to OD_600_ = 5 and harvested by centrifugation at 4,700 x *g* for 10 minutes. The cell pellet was resuspended in 200 μL of cold TBS supplemented with 1 mM PMSF and lysed via three cycles of bead beating (30-second durations with 30-second intervals on ice; BioSpec Products). Cell lysates were centrifuged at 4,700 x *g* to remove any unbroken cells, and subjected to SDS-PAGE and immunoblot analysis.

### SDS-PAGE and immunoblot analysis

Cell lysates containing His-tagged proteins or purified protein samples were mixed with Laemmli buffer and analyzed by Tris-glycine SDS-PAGE with 12% Tris-HCl gels at 160 V for 55 minutes. With the exception of the heat-modifiable gel shift analysis, all samples were heated at 95 °C for 10 minutes. Gels were then visualized by InstantBlue Coomassie Protein Stain (Abcam) or subjected to immunoblotting analysis. For the latter, proteins on the gel were transferred onto a polyvinylidene fluoride membrane (Bio-Rad) using a Transblot SD semi-dry transfer system (Bio-Rad) at 25 V for 30 minutes. The membrane was then blocked for at least 1 hour in 1x casein blocking buffer (Sigma–Aldrich). To detect His-tagged proteins, the membrane was incubated for 1 hour with monoclonal α-His antibodies conjugated to the horseradish peroxidase (1:5000 dilution; Qiagen). Blots were visualized with the addition of Lumina Forte Western HRP Substrate (MilliporeSigma) to the membrane.

### SEC-MALS analysis

Prior to SEC-MALS analysis, a calibration run with monomeric bovine serum albumin (BSA; Sigma-Aldrich) was performed, and the settings (calibration constant for TREOS detector; Wyatt Technology) that gave the exact molar mass of BSA (66.4 kDa) were used for subsequent molar mass calculations. MSMEG_0317 and truncated variants (purified and concentrated) were injected into a Superdex 200 Increase 10/300 GL column pre-equilibrated with TBS-D. Light scattering and refractive index (*n*) data were collected online using miniDAWN TREOS (Wyatt Technology) and Optilab T-rEX (Wyatt Technology), respectively, and analyzed bby ASTRA 6.1.5.22 software (Wyatt Technology). Protein-conjugate analysis in the ASTRA software was employed to calculate nonproteinaceous components of the complex (i.e., DDM). The refractive index increment *dn/dc* values (where c is the sample concentration) of 0.143 ml/g and 0.185 ml/g were used for DDM and protein complex respectively as previously reported (26).

### Lipid profiling

cKO cells grown in the presence or absence of acetamide normalized to OD_600_ = 1 were incubated for 2 hours at 37 °C in 7H9 media containing [^14^C]-acetate (final concentration 0.5 μCi/ml; Perkin Elmer). Labeled cells were harvested by centrifugation at 5,000 x *g* for 3 minutes, and cell pellet was resuspended in chloroform-methanol (2:1) solution. The suspension was subjected to three cycles of sonication (30 seconds duration) and brief vortexing between each cycle. Methanol and ultrapure water were further added to give a biphasic chloroform-methanol-water (1:1:0.8) mixture, and centrifuged to achieve phase separation. The organic phase (chloroform, contains extractable lipids) was collected and air-dried overnight in a fumehood. The cell debris in the aqueous phase, containing covalently-linked MAs, was dried in a speed-vac Concentrator Plus (Eppendorf). MAs were liberated from the dried cell debris by heating at 90 °C in 40% (w/v) tetrabutylammonium hydroxide (TBAH; Sigma-Aldrich) for 2 hours. After cooling, the mixture was acidified, extracted with *n*-hexane, and air-dried overnight in a fumehood (27). Extractable lipids and liberated MAs were analyzed via TLC (TLC Silica gel 60 F_254_; Merck Millipore) developed in chloroform-methanol-water (30:8:1), and visualized by phosphor imaging (GE Healthcare).

## Supporting information

This article contains supporting information.

## Acknowledgments

We thank Ross Tomaino (Taplin MS facility, Harvard Medical School) for help with MS/MS and protein identification. We also thank Ruby Hao Sun for sharing her data with us, and Dr. Sizhun Li for constructing the pJEB402-hyg plasmid used in this study.

## Funding and additional information

This work was supported by the Singapore of Ministry of Education Academic Research Fund Tier 2 grants (MOE2018-T2-1-128 and MOE000116, to S.-S.C.).

## References

1. Brennan, P. J., and Nikaido, H. (1995) The envelope of mycobacteria. Annu. Rev. Biochem. 64, 29–63

2. Jackson, M. (2014) The mycobacterial cell envelope-lipids. Cold Spring Harb. Perspect. Med. 4, a021105

3. Bansal-Mutalik, R., and Nikaido, H. (2014) Mycobacterial outer membrane is a lipid bilayer and the inner membrane is unusually rich in diacyl phosphatidylinositol dimannosides. Proc. Natl. Acad. Sci. U.S.A.111, 4958

4. Banerjee, A., Dubnau, E., Quemard, A., Balasubramanian, V., Um, K. S., Wilson, T., Collins, D., de Lisle, G., and Jacobs, W. R. (1994) inhA, a Gene Encoding a Target for Isoniazid and Ethionamide in Mycobacterium tuberculosis. Science 263, 227–230

5. Xu, Z., Meshcheryakov, V. A., Poce, G., and Chng, S.-S. (2017) MmpL3 is the flippase for mycolic acids in mycobacteria. Proc. Natl. Acad. Sci. U.S.A. 114, 7993

6. Belisle, J. T., Vissa, V. D., Sievert, T., Takayama, K., Brennan, P. J., and Besra, G. S. (1997) Role of the Major Antigen of Mycobacterium tuberculosis in Cell Wall Biogenesis. Science 276, 1420–1422

7. Rainczuk, A. K., Klatt, S., Yamaryo-Botté, Y., Brammananth, R., McConville, M. J., Coppel, R. L., and Crellin, P. K. (2020) MtrP, a putative methyltransferase in Corynebacteria, is required for optimal membrane transport of trehalose mycolates. J. Biol. Chem. 295, 6108–6119

8. Cashmore, T. J., Klatt, S., Brammananth, R., Rainczuk, A. K., Crellin, P. K., McConville, M. J., and Coppel, R. L. (2021) MmpA, a Conserved Membrane Protein Required for Efficient Surface Transport of Trehalose Lipids in Corynebacterineae. Biomolecules 11, 1760

9. Yamaryo-Botte, Y., Rainczuk, A. K., Lea-Smith, D. J., Brammananth, R., van der Peet, P. L., Meikle, P., Ralton, J. E., Rupasinghe, T. W. T., Williams, S. J., Coppel, R. L., Crellin, P. K., and McConville, M. J. (2015) Acetylation of Trehalose Mycolates Is Required for Efficient MmpL-Mediated Membrane Transport in Corynebacterineae. ACS Chem. Biol. 10, 734–746

10. Collins, M. D., Goodfellow, M., and Minnikin, D. E. (1982) A Survey of the Structures of Mycolic Acids in Corynebacterium and Related Taxa. Microbiology 128, 129–149

11. Klatt, S., Brammananth, R., O’Callaghan, S., Kouremenos, K. A., Tull, D., Crellin, P. K., Coppel, R. L., and McConville, M. J. (2018) Identification of novel lipid modifications and intermembrane dynamics in Corynebacterium glutamicum using high-resolution mass spectrometry. J. Lipid Res. 59, 1190–1204

12. Belardinelli, J. M., Yazidi, A., Yang, L., Fabre, L., Li, W., Jacques, B., Angala, S. K., Rouiller, I., Zgurskaya, H. I., Sygusch, J., and Jackson, M. (2016) Structure-Function Profile of MmpL3, the Essential Mycolic Acid Transporter from Mycobacterium tuberculosis. ACS Infect. Dis. 2, 702–713

13. Cashmore, T. J., Klatt, S., Yamaryo-Botte, Y., Brammananth, R., Rainczuk, A. K., McConville, M. J., Crellin, P. K., and Coppel, R. L. (2017) Identification of a Membrane Protein Required for Lipomannan Maturation and Lipoarabinomannan Synthesis in Corynebacterineae*. J. Biol. Chem. 292, 4976–4986

14. Sassetti, C. M., Boyd, D. H., and Rubin, E. J. (2003) Genes required for mycobacterial growth defined by high density mutagenesis. Mol. Microbiol. 48, 77–84

15. Zhang, Y. J., Ioerger, T. R., Huttenhower, C., Long, J. E., Sassetti, C. M., Sacchettini, J. C., and Rubin, E. J. (2012) Global Assessment of Genomic Regions Required for Growth in Mycobacterium tuberculosis. PLOS Pathog. 8, e1002946

16. Fukuda, T., Matsumura, T., Ato, M., Hamasaki, M., Nishiuchi, Y., Murakami, Y., Maeda, Y., Yoshimori, T., Matsumoto, S., Kobayashi, K., Kinoshita, T., and Morita, Y. S. (2013) Critical roles for lipomannan and lipoarabinomannan in cell wall integrity of mycobacteria and pathogenesis of tuberculosis. mBio 4, e00472

17. Banerjee, R., Vats, P., Dahale, S., Kasibhatla, S. M., and Joshi, R. (2011) Comparative Genomics of Cell Envelope Components in Mycobacteria. PLOS ONE 6, e19280

18. Belardinelli, J. M., Stevens, C. M., Li, W., Tan, Y. Z., Jones, V., Mancia, F., Zgurskaya, H. I., and Jackson, M. (2019) The MmpL3 interactome reveals a complex crosstalk between cell envelope biosynthesis and cell elongation and division in mycobacteria. Sci. Rep. 9, 10728

19. Patel, O., Brammananth, R., Dai, W., Panjikar, S., Coppel, R. L., Lucet, I. S., and Crellin, P. K. (2022) Crystal structure of the putative cell-wall lipoglycan biosynthesis protein LmcA from Mycobacterium smegmatis. Acta Crystallogr. D 78, 494–508

20. Kavunja, H. W., Biegas, K. J., Banahene, N., Stewart, J. A., Piligian, B. F., Groenevelt, J. M., Sein, C. E., Morita, Y. S., Niederweis, M., Siegrist, M. S., and Swarts, B. M. (2020) Photoactivatable Glycolipid Probes for Identifying Mycolate-Protein Interactions in Live Mycobacteria. J. Am. Chem. Soc. 142, 7725–7731

21. Su, C.-C., Klenotic, P. A., Cui, M., Lyu, M., Morgan, C. E., and Yu, E. W. (2021) Structures of the mycobacterial membrane protein MmpL3 reveal its mechanism of lipid transport. PLOS Biol. 19, e3001370

22. Zhang, B., Li, J., Yang, X., Wu, L., Zhang, J., Yang, Y., Zhao, Y., Zhang, L., Yang, X., Yang, X., Cheng, X., Liu, Z., Jiang, B., Jiang, H., Guddat, L. W., Yang, H., and Rao, Z. (2019) Crystal Structures of Membrane Transporter MmpL3, an Anti-TB Drug Target. Cell 176, 636-648.e613

23. Gupta, K. R., Gwin, C. M., Rahlwes, K. C., Biegas, K. J., Wang, C., Park, J. H., Liu, J., Swarts, B. M., Morita, Y. S., and Rego, E. H. (2022) An essential periplasmic protein coordinates lipid trafficking and is required for asymmetric polar growth in mycobacteria. eLife 11, e80395

24. Organization, W. H. (2022) Global Tuberculosis Report 2022.

25. Parish, T., and Stoker, N. G. (2000) Use of a flexible cassette method to generate a double unmarked Mycobacterium tuberculosis tlyA plcABC mutant by gene replacement. Microbiology 146, 1969–1975

26. Yeow, J., Tan, K. W., Holdbrook, D. A., Chong, Z. S., Marzinek, J. K., Bond, P. J., and Chng, S. S. (2018) The architecture of the OmpC-MlaA complex sheds light on the maintenance of outer membrane lipid asymmetry in Escherichia coli. J. Biol. Chem. 293, 11325–11340

27. Hsu, F. F., Soehl, K., Turk, J., and Haas, A. (2011) Characterization of mycolic acids from the pathogen Rhodococcus equi by tandem mass spectrometry with electrospray ionization. Anal. Biochem. 409, 112–122

