## Supplementary information for "Defining a critical role of an essential membrane protein in mycolic acid transport in mycobacteria"

**This file contains:**

**Supplementary Tables S1 and S2.**

**Supplementary Figures S1 and S2.**

### Supplementary tables

#### Table S1: Plasmids used in this study.

| Plasmids | Description | References |
|---|---|---|
| pYUB854 | Used for homologous recombination in mycobacteria, contains an origin of replication for *E. coli* (*oriE*) and λ-cos sites; $Hyg^R$ | (1) |
| pYUB854-0317-1CO | pYUB854 containing a *spec*$^R$ cassette flanked by ~1000 bases of the 5'- and 3'-untranslated regions of the *msmeg_0317* gene | This study |
| pMyC-kan | Mycobacterial expression vector with an acetamidase promoter, contains an origin of replication for *M. smegmatis* (*oriM*) and an *oriE*; $Kan^R$ | (2) |
| pMyC-0317 | pMyC-kan; encodes full length MSMEG_0317; $Kan^R$ | This study |
| pJEB402-kan | Mycobacterial chromosomal integration vector, contains a constitutive MOP promoter and L5 *attB* site; $Kan^R$ | (3) |
| pJEB402-kan-N0317 | pJEB402-kan; encodes full length MSMEG_0317 with a N-terminal $His_6$ tag; $Kan^R$ | This study |
| pJEB402-kan-C0317 | pJEB402-kan; encodes full length MSMEG_0317 with a C-terminal $His_8$ tag; $Kan^R$ | This study |
| pET28b(+) | *E. coli* expression vector with a T7-lac inducible promoter; $Kan^R$ | Novagen |
| pET22/42 | *E. coli* expression vector with a T7-lac inducible promoter, contains multiple cloning sites of pET42a(+) in pET22b(+) backbone; $Amp^R$ | Lab collection |
| pET28b-N0317 | pET28b(+); encodes full length MSMEG_0317 with a N-terminal $His_6$ tag; $Kan^R$ | This study |
| pET22/42-C0317 | pET22/42; encodes full length MSMEG_0317 with a C-terminal $His_8$ tag; $Amp^R$ | This study |
| pET28b-N0317ΔCD | pET28b; encodes 0317ΔCD ($MSMEG\_0317_{1-350}$) with a N-terminal $His_6$ tag; $Kan^R$ | This study |
| pET22/42-C0317ΔNH | pET22/42; encodes 0317ΔNH ($MSMEG\_0317_{33-398}$) with a C-terminal $His_8$ tag; $Amp^R$ | This study |
| pET28b-N0317ΔCHD | pET28b; encodes 0317ΔCHD ($MSMEG\_0317_{1-323}$) with a N-terminal $His_6$ tag; $Kan^R$ | This study |
| pET22/42-S-C0317 | pET22/42; encodes S-0317 ($MSMEG\_0317_{33-323}$) with a C-terminal $His_8$ tag; $Amp^R$ | This study |
| pJEB402-hyg | pJEB402-kan with the $Kan^R$ cassette replaced with a $Hyg^R$ cassette | This study |
| pJEB402-hyg-Rv0227c | pJEB402-hyg; encodes full length Rv0227c; $Hyg^R$ | This study |
| pJEB402-hyg-0317 | pJEB402-hyg; encodes full length MSMEG_0317; $Hyg^R$ | This study |

| | | |
|---|---|---|
| pJEB402-hyg-0317ΔNH | pJEB402-hyg; encodes 0317ΔNH (MSMEG_0317$_{33\text{-}398}$); Hyg$^R$ | This study |
| pJEB402-hyg-0317ΔCHD | pJEB402-hyg; encodes 0317ΔCHD (MSMEG_0317$_{1\text{-}323}$); Hyg$^R$ | This study |
| pJEB402-hyg-0317ΔCD | pJEB402-hyg; encodes 0317ΔCD (MSMEG_0317$_{1\text{-}350}$); Hyg$^R$ | This study |
| pJEB402-hyg-C0317 | pJEB402-hyg; encodes full length MSMEG_0317 with a C-terminal His$_8$ tag; Kan$^R$ | This study |
| pJEB402-hyg-C0317ΔNH | pJEB402-hyg; encodes 0317ΔNH (MSMEG_0317$_{33\text{-}398}$) with a C-terminal His$_8$ tag; Hyg$^R$ | This study |
| pJEB402-hyg-C0317ΔCHD | pJEB402-hyg; encodes 0317ΔCHD (MSMEG_0317$_{1\text{-}323}$) with a C-terminal His$_8$ tag; Hyg$^R$ | This study |
| pJEB402-hyg-C0317ΔCD | pJEB402-hyg; encodes 0317ΔCD (MSMEG_0317$_{1\text{-}350}$) with a C-terminal His$_8$ tag; Hyg$^R$ | This study |

**Table S2: Primers used in this study.**

| Primers | Sequence (5' to 3') |
|---|---|
| pYUB854-5UTR FP | ATATACTAGTGCGGTCTTGAGCATGACCTG |
| pYUB854-5UTR RP | ATCAAAGCTTACCGGGCCTCCTTCTGTGT |
| pYUB854-3UTR FP | ATATAAGCTTGAACTGCGGTCCTGCTGC |
| pYUB854-3UTR RP | TATAGCTAGCGTCTATCAGCGCCTCCACC |
| pMyC-0317 FP | AAGGGAGTCCACATGGCCCGGACGCGGTCGATGCCG |
| pMyC-0317 RP | GTGGTGGTGGTGCGATCAGATCGGTCGGTCCGGTGGCAGA |
| pJEB402-hyg-Rv0227c FP | ATATCATATGTTGCGGTTCGCCGCGTGCGGC |
| pJEB402-hyg-Rv0227c RP | ATATAAGCTTAGCGATGATTAGCGCCGCTCGGG |
| pJEB402-kan-N/C0317 FP | TCGAGAATTCTTAACTTTAAGAAGGAGATATAC |
| pJEB402-kan-N/C0317 RP | ACGTAAGCTTGCAGCCTAGGTATTAATCAATTAGTGGTG |
| pET28b-N0317 FP | AATTCATATGAAGGGCAAGATCGCCAAGATC |
| pET28b-N0317 RP | ATATAAGCTTCTACCGTGTCCACAGCGCGATG |
| pET22/42-C0317 FP | AATTCATATGAACCGCGCTGTGGCG |
| pET22/42-C0317 RP | ATATCTCGAGGATCGGTCGGTCCGGTGGCAGAT |
| pET28b-N0317ΔCD FP | CGCACCTGAGAAGCTTGCGGCCGCA |
| pET28b-N0317ΔCD RP | TTCTCAGGTGCGCAGCGCGAACGA |
| pET28b-N0317 ΔCHD FP | ACACGGTGAGAAGCTTGCGGCCGCA |
| pET28b-N0317 ΔCHD RP | TTCTCACGTGTCCACAGCGCGAT |
| pET22/42-C0317 ΔNH FP | CATATGACCACCTACACCAAGGGC |
| pET22/42-C0317 ΔNH RP | GGTGGTCATATGTATATCTCCTTCTTAAAG |
| pET22/42-S-C0317 FP | AATTCATATGAAGGGCAAGATCGCCA |
| pET22/42-S-C0317 RP | ATATAAGCTTCTACCGTGTCCACAGCG |
| pJEB402-hyg-0317 FP | AATTCATATGAACCGCGCTGTGGCG |
| pJEB402-hyg-0317 RP | ATATAAGCTTCAGGACCGCAGTTCTCAGATC |
| pJEB402-hyg-0317ΔNH FP | AATTCATATGAAGGGCAAGATCGCCA |
| pJEB402-hyg-0317ΔNH RP | ATATAAGCTTCAGGACCGCAGTTCTCAGATC |
| pJEB402-hyg-0317ΔCD FP | AATTCATATGAACCGCGCTGTGGCG |
| pJEB402-hyg-0317ΔCD RP | ATATAAGCTTTCAGGTGCGCAGCGCGAACGA |
| pJEB402-hyg-0317ΔCHD FP | AATTCATATGAACCGCGCTGTGGCG |
| pJEB402-hyg-0317ΔCHD RP | CGAGTGCGGCCGAAGCTTCTCA |
| pJEB402-hyg-C0317 FP | ATATGAGATATACATATGAACCGCGCTGTG |
| pJEB402-hyg-C0317 RP | ACGTAAGCTTGCAGCCTAGGTATTAATCAATTAGTGGTG |
| pJEB402-hyg-C0317ΔNH FP | GAAGGAGATATACATATGAAGGGCAAGATCGCCAAG |
| pJEB402-hyg-C0317ΔNH RP | GTCGATGTCGAGCGGGATCTTGGCGATCTTGCCCTT |
| pJEB402-hyg-C0317ΔCHD FP | ATCGCGCTGTGGACACGGCACCACCACCACCACCAC |
| pJEB402-hyg-C0317ΔCHD RP | GTCGATGTCGAGCGGGATCTTGGCGATCTTGCCCTT |
| pJEB402-hyg-C0317ΔCD FP | TCGTTCGCGCTGCGCACCCACCACCACCACCACCAC |
| pJEB402-hyg-C0317ΔCD RP | ATTAATCAATTAGTGGTGGTGGTGGTGGTGGTGGTG |

*Sites for restriction enzyme cleavage, where relevant, are underlined.

### Supplementary figures

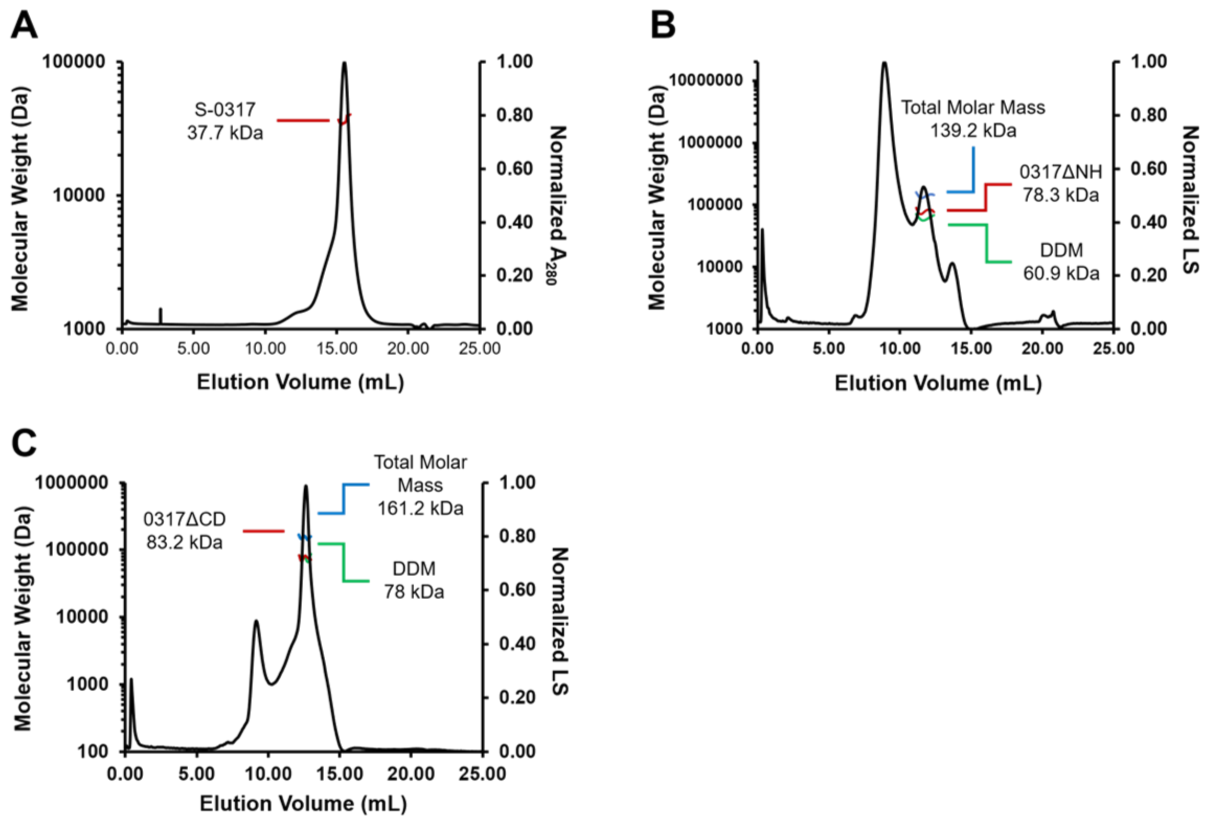


**Figure S1**: **SEC-MALS analysis of MSMEG_0317 truncated variants. (A)** 0317ΔCD exists as a dimer. Total molar mass: 161.2 (± 5.4 %) kDa; modifier (DDM) molar mass: 78.0 (± 15.7 %); protein molar mass: 83.2 (± 4.4 %) kDa (measured), 72.5 kDa (predicted for 0317ΔCD dimer). **(B)** 0317ΔNH exists as a dimer. Total molar mass: 139.2 (± 4.9 %) kDa; modifier (DDM) molar mass: 60.9 (± 12.2 %); protein molar mass: 78.3 (± 4.4 %) kDa (measured), 82.6 kDa (predicted for 0317ΔNH dimer). **(C)** S-0317 exists as a monomer. Protein molar mass: 37.7 (± 8.6 %) kDa (measured), 33.5 kDa (predicted for S-0317 monomer). Statistical analysis errors are reported as percentages.

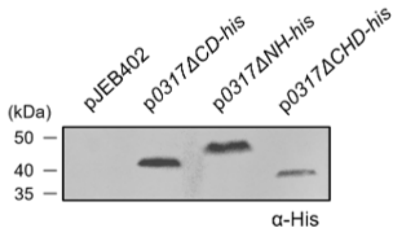


**Figure S2: Truncated MSMEG_0317 variants are expressed *in vivo*.** Immunoblot analysis of cKO cell lysates expressing His-tagged 0317ΔCD, 0317ΔNH, and 0317ΔCHD, under the constitutive MOP promoter.

### References

1. Bardarov, S., Bardarov, S., Pavelka, M. S., Sambandamurthy, V., Larsen, M., Tufariello, J., Chan, J., Hatfull, G., and Jacobs, W. R. (2002) Specialized transduction: an efficient method for generating marked and unmarked targeted gene disruptions in *Mycobacterium tuberculosis*, *M. bovis BCG* and *M. smegmatis*. *Microbiology* **148**, 3007-3017
2. Beckham, K. S. H., Staack, S., Wilmanns, M., and Parret, A. H. A. (2020) The pMy vector series: A versatile cloning platform for the recombinant production of mycobacterial proteins in *Mycobacterium smegmatis*. *Protein Sci.* **29**, 2528-2537
3. Lee, M. H., Pascopella, L., Jacobs, W. R., Jr., and Hatfull, G. F. (1991) Site-specific integration of mycobacteriophage L5: integration-proficient vectors for *Mycobacterium smegmatis*, *Mycobacterium tuberculosis*, and bacille Calmette-Guérin. *Proc. Natl. Acad. Sci. U.S.A.* **88**, 3111-3115
